# Electroencephalographic dynamics of rhythmic breath-based meditation

**DOI:** 10.1101/2022.03.09.483685

**Authors:** Vaibhav Tripathi, Lakshmi Bhasker, Chhaya Kharya, Manvir Bhatia, Vinod Kochupillai

## Abstract

Meditation has been practised for millennia but the neuroscientific understanding of the dynamics is still lacking. Sudarshan Kriya Yoga (SKY) is an evidence based breathing based meditation technique that utilizes rhythmic breathing to induce a deep state of relaxation and calm. Multiple studies have found benefits of the SKY technique from genetic, physiological, psychological to behaviour levels. We collected Electroencephalographic (EEG) data in 43 subjects who underwent the SKY technique and analysed the brain rhythms at different stages of the technique namely preparatory breathing (Pranayama), rhythmic breathing (Kriya) and meditation (Yoga Nidra) using newly developed methods to analyse periodic and aperiodic components. Alpha waves amplitude in the parieto-occipital region decreased as the rhythmic breathing progressed and dropped sharply during the meditation period. Theta amplitudes and peak frequency increased in the centro-frontal region during the rhythmic breathing period but were marked by sustained low theta waves during the meditation period. The delta wave amplitude was not affected by breathing but both delta band power and peak frequency increased during the meditation period in the centro-frontal region. We also saw a decrease in the 1/f aperiodic signal across the brain during the meditation period suggesting a modification of excitation-inhibition balance. We see an overall slowing down of brain oscillations from alpha to theta to delta as the meditation progressed. The paper studies in depth the transitional dynamics of the SKY technique analysing the alpha, theta, delta waves and aperiodic signals and demonstrates that each phase in a breathing based meditation has a unique electrophysiological signature.

## Introduction

Meditation has been a mainstay of spiritual practices since millenia. Multiple studies have shown the benefits of meditation to boost mental and physical health^1–3^. The development of Electroencephalography (EEG), functional MRI (fMRI) has further the research on understanding the spatiotemporal dynamics of meditation^4–8^.

Respiration has been linked to modulate electrical and oscillatory activity in the brain^9–14^. Breathing practices has been an integral part of the Yoga system and have been practised for millenia for improved mental and physical well being. Recent studies have shown the effect of breathing on mental states and electrical activity in the brain^15–17^.

The Yoga Sutras of Patanjali, the de-facto guide to Yoga holds breathing techniques (Pranayama) as one of the limbs of Yoga which allows better transition to deeper meditation states^18^. Rhythmic breathing techniques like the Sudarshan Kriya Yoga (SKY) utilizes the breath to induce deep meditation states. Studies have shown the effects of SKY on anxiety and stress reduction^19^, increase in natural killer cells^20^ and blood lactate decrease with antioxidant increase^20^. SKY has shown to reduce symptoms of Post Traumatic Stress Disorder (PTSD) in war veterans^21^ and even act as an antidepressant^22,23^. Randomized controlled trials have shown that SKY technique is better at enhancing social connect, and mental well being as compared to other mindfulness practices which do not include the breathing component^24,25^. SKY has shown to induce immediate effects on the gene expression profile in immune cells^26^ and may improve immunity, reduce aging and decrease stress through transcriptional regulation^27^.

Previous studies have attempted to determine the effects of SKY on the brain waves as detected using EEG. SKY induces a state of wakeful alertness^22^, and enhances theta band activity and coherence^28^ and inter-hemispheric synchronization of the brain rhythms^29^. The analysis of these studies were limited to differences in the oscillatory activity before and after the SKY practice. In the current paper, we look at the oscillatory dynamics during the different stages of SKY meditation practice namely preparatory breathing (Pranayama), rhythmic breathing (Kriya) and meditation (Yoga Nidra) along with comparing with resting state EEG before and after the practice. We found that the theta waves increased in both amplitude and frequency during the breathing phase in the centro-frontal regions. During the resting phase in supine position, we saw increase in theta waves in the centro-frontal regions and decrease in the alpha amplitudes in the parieto-occipital regions. The yoga nidra period was dominated by delta and theta rhythms. We also analysed the aperiodic signals which have been demonstrated to be a marker of excitation-inhibition balance^30–32^ and found that the exponent of the aperiodic signal decreased across the brain during the meditation period. The results depict transition of the brain from awake and active represented by alpha band activity to a calm, relax meditative state with high delta and theta which illustrates a state between sleep and awake, something similar to the state of “Turiya” as mentioned in ancient yogic literature. Ours is one of the first studies to look at the spatiotemporal dynamics of brain waves during different phases of a rhythmic breath-based meditation.

## Results

In order to analyse the data across the different parts of the SKY meditation, we divided the duration into five events: resting-pre, pranayama, kriya, yoga nidra and resting-post (Figure 1a). We extracted the peak frequency, amplitude and band power component (described in the methods section) for each event (as given in Figure 1b). We then compared the components in the alpha (7-13 Hz), theta (4-7 Hz) and delta (1-4 Hz) bands and aperiodic signals across the five events.

**Figure 1.**
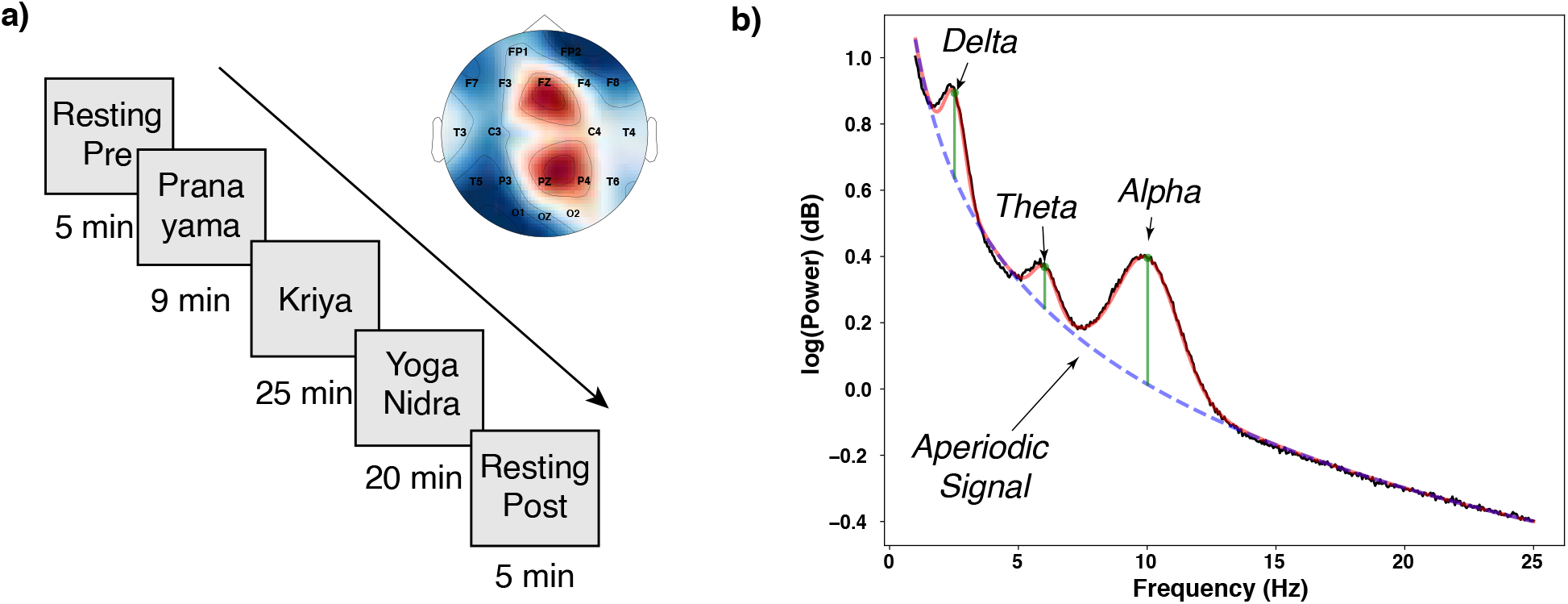
Protocol and Analysis. a) Protocol for the breath-based meditation. EEG was recorded in a 10/20 arrangement from participants undergoing the various steps in the Sudarshan Kriya Yoga (SKY) meditation which involved breathing exercises called pranayama for 18 minutes (the first 9 minutes analysed in the paper) followed by the rhythming breathing (Kriya) followed by rest in the supine position. Pre and post the meditation, we recorded resting state activity. b) The data was cleaned, preprocessed and frequency spectrum was plotted for a time window of 4 s with step size of 1 s and extracted the spectral properties including the delta, theta and alpha peak frequency, amplitude and band power werealong with aperiodic signal characteristics using FOOOF toolbox. The diagram shows relevant features extracted from the frequency spectrum.

Figure 2 shows the alpha band power, peak frequency and amplitude across the five events. We performed repeated measures ANOVA across the five events for all subjects for each individual channel separately for band power, frequency and amplitude and corrected for multiple comparisons across channels using bonferroni correction. We see a statistical significant decrease of alpha peak amplitude in the parieto-temporal and occipital regions as the SKY meditation progressed (P4: F(4,164)=36.85, p < 0.00001, Post hoc analysis using Tukey’s test found differences between Yoga Nidra (M=5.13, SD=1.24), Post (M=5.19, SD=1.66) and Pre (M=2.38, SD=0.33), Pranayama (M=2.36, SD=0.40), SKY (M=2.51, SD=0.30, no other pairs had significant differences. Similar statistics were computed for P3, PZ, T5, T6, O1 and OZ channels). We see a gradual decrease in the peak amplitude during the yoga nidra period as can be seen from last panel in Fig. 2c. The peak frequency did not change across the events (P4: F(4,164)=0.95, p=0.43). The band power increased significantly in parieto-temporal and frontal regions (F(4,164)=10.21, p<0.00001 Tukey’s post hoc differences were found between Yoga Nidra (M=2.66, SD=0.28) and Pre (M=2.38, SD=0.33), Pranayama (M=2.36, SD=0.40), no other pairs had significant differences. We found similar statistics for FP1, F3, FZ, T3, T5, T6 and P3 channels).

**Figure 2.**
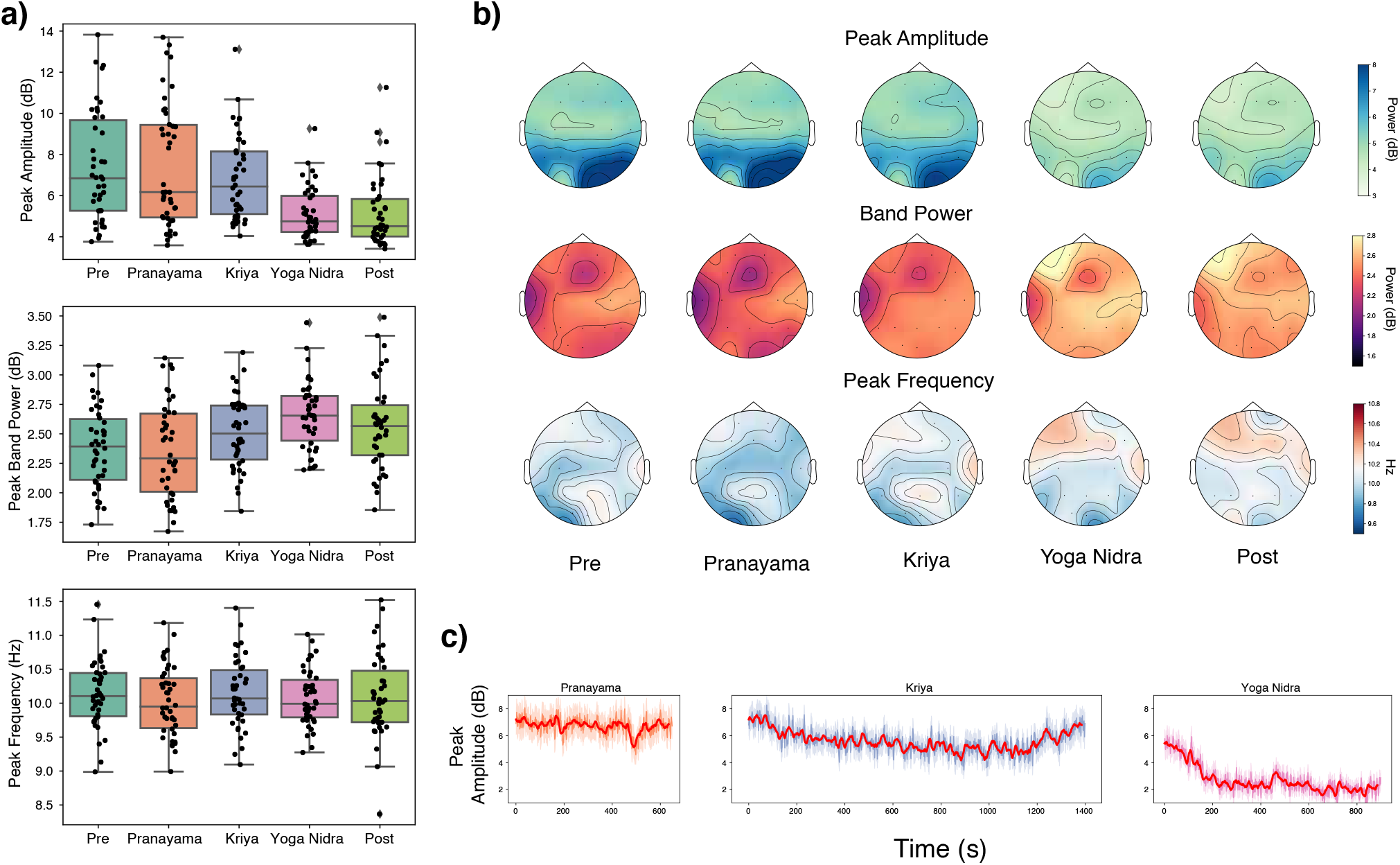
Effect on Alpha wave dynamics: a) Bar plots show the peak amplitude, band power and frequency for the alpha band computed using the FOOOF toolbox (Donogue et. al. 2020) for P4 electrode. b) Topomap plots for the peak amplitude, band power and peak frequency for the alpha waves across the various parts of the SKY meditation. c) Change of peak amplitude across the meditation, we see a slow decrease in the alpha amplitude during the Kriya period which sharply drops during the Yoga Nidra period.

We then analysed the theta band power, peak frequency and amplitude across the five events (as shown in figure 3) and found that the theta peak amplitude increased significantly during the kriya period and decreased afterwards in the frontal, temporal and parietal regions (FZ channel: F(4,164)=6.49, p<0.00007, Tukey’s post hoc differences were found between SKY (M=4.75, SD=1.01) and Pre (M=4.10, SD=0.89), Pranayama (M=3.98, SD=1.21) and Post (M=3.93, SD=1.10). Channels F4, F8, T4, PZ, P4 showed similar behavior) whereas the band power increased during the yoga nidra period in the temporal-parietal regions (T4: F(4,164)=12.95, p<0.00001, Tukey’s post hoc differences were found between Yoga Nidra (M=2.67, SD=0.29), Post (M=2.51, SD=0.59) and Pre (M=2.26, SD=0.45), Pranayama (M=2.19, SD=0.50) and SKY (M=2.16, SD=0.36), no other differences were found. We found similar behavior for channels T5, T6, P3, P4, PZ). The theta peak frequency increased till the kriya period and then decreased during the post resting phase across centro-temporal and parietal regions (C4 Channel: F(4,164)=7.88, p<0.00001 with Tukey’s post hoc differences between Post (M=5.03, SD=1.42) and Pre (M=5.58, SD=0.24), Pranayama (M=5.64, SD=0.28), SKY (M=5.76, SD=0.22). And we found similar statistics for channels C3, T4, T5, T6, P3, P4, PZ). Panel 3C shows sustained increase in theta peak amplitude during the kriya period followed by elevated but reduced activity during the yoga nidra phase.

**Figure 3.**
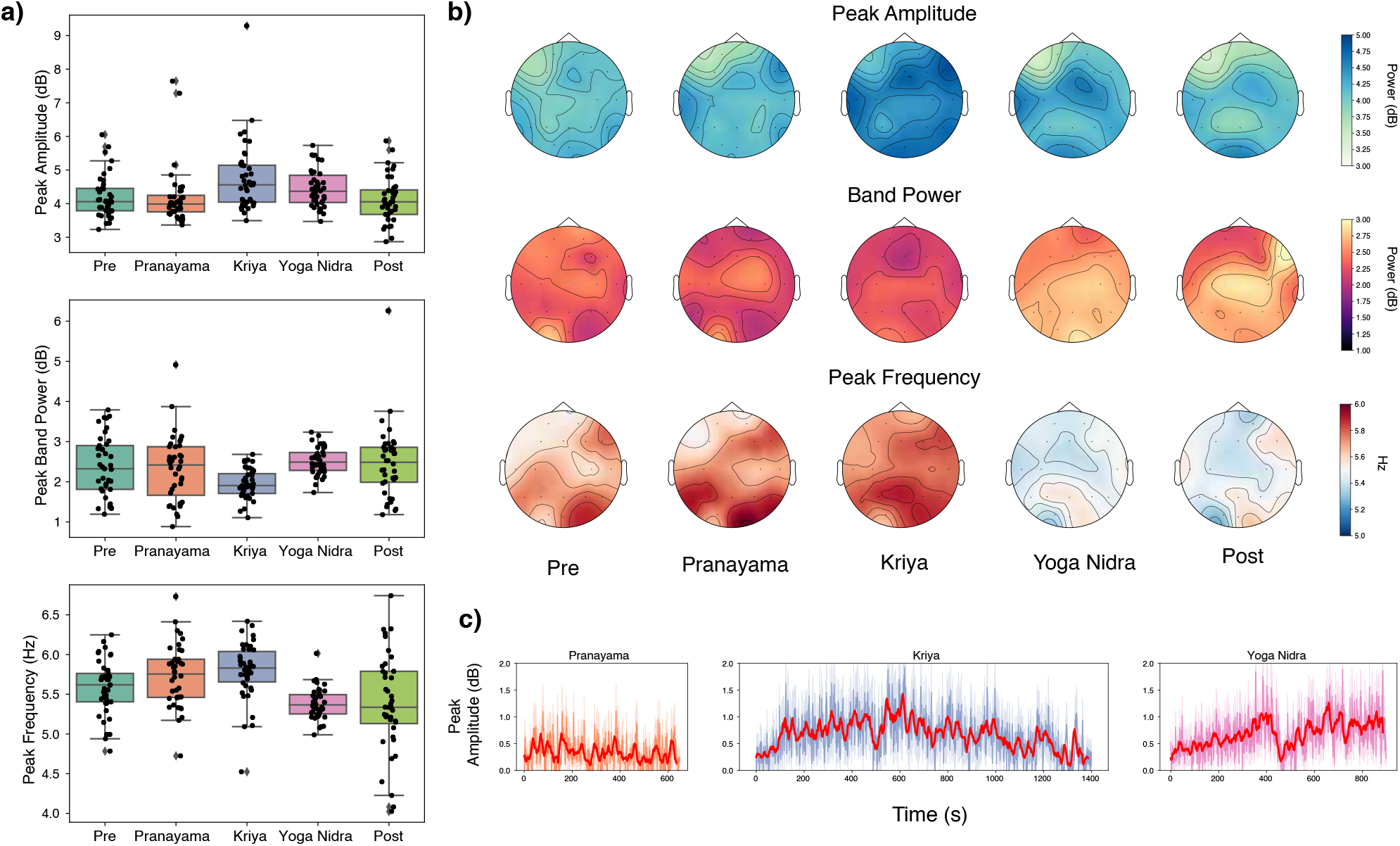
Effect on Theta wave dynamics: a) Bar plots show the peak amplitude, band power and frequency for the theta band computed using the FOOOF toolbox (Donogue et. al. 2020) for FZ electrode. b) Topomap plots for the peak amplitude, band power and peak frequency for the theta waves across the various parts of the SKY meditation. c) Change of peak amplitude across the meditation, we see a theta increase during both the Kriya and the Yoga Nidra periods of the practice.

Looking at the delta band components, we find that delta band power increased significantly in the frontal (FZ channel: F(4,164)=25.11, p<0.00001, we found Tukey’s post hoc differences between Yoga Nidra (M=2.66, SD=0.30), Post (M=2.53, SD=0.75) and Pre (M=2.02, SD=0.54), Pranayama (M=1.99, SD=0.63), SKY (M=1.93, SD=0.37), FP1, FP2,F3, F4 had similar statistics) and left centro-temporal regions (F(4,164)=9.79, p<0.00001, Tukey’s post hoc differences were found between Yoga Nidra (M=2.78, SD=0.30), Post (M=2.56, SD=1.30) and Pre (M=2.13, SD=0.56), Pranayama (M=2.08, SD=0.58), SKY (M=2.06, SD=0.45). Panel 4c depicts a sharp rise in delta amplitude during the yoga nidra period. The delta peak frequency increased across the whole brain during the yoga nidra period (FZ Channel: F(4,164)=27.39, p<0.00001, Tukey’s post hoc differences were found between Yoga Nidra (M=2.34, SD=0.17), Post (M=2.27, SD=0.40) and Pre (M=1.96, SD=0.26), Pranayama (M=1.88, SD=0.23), SKY (M=2.06, SD=0.24)).

Recent literature on aperiodic signals^30,32,33^ have found association of the exponent of the aperiodic signals with age related decline^32^, states of conciousness^34^, attention and working memory^31^ and even meditation^35^. As seen from Fig. 5c, we see a sharp decrease in the aperiodic exponent in the frontal and left temporal regions during the yoga nidra and post resting period (FZ Channel: F(4,164)=29.82, p<0.00001, Tukey’s post hoc differences between Yoga Nidra (M=7.26, SD=1.30), Post (M=7.31, SD=1.54) and Pre (M=8.91, SD=1.88), Pranayama (M=8.74, SD=2.48), SKY (M=9.01, SD=1.77). We find similar statistics for channels FP1, FP2, F4, F8 and T3). Ours is the first study to look at the effect of breathing based meditation on the aperiodic signals.

**Figure 4.**
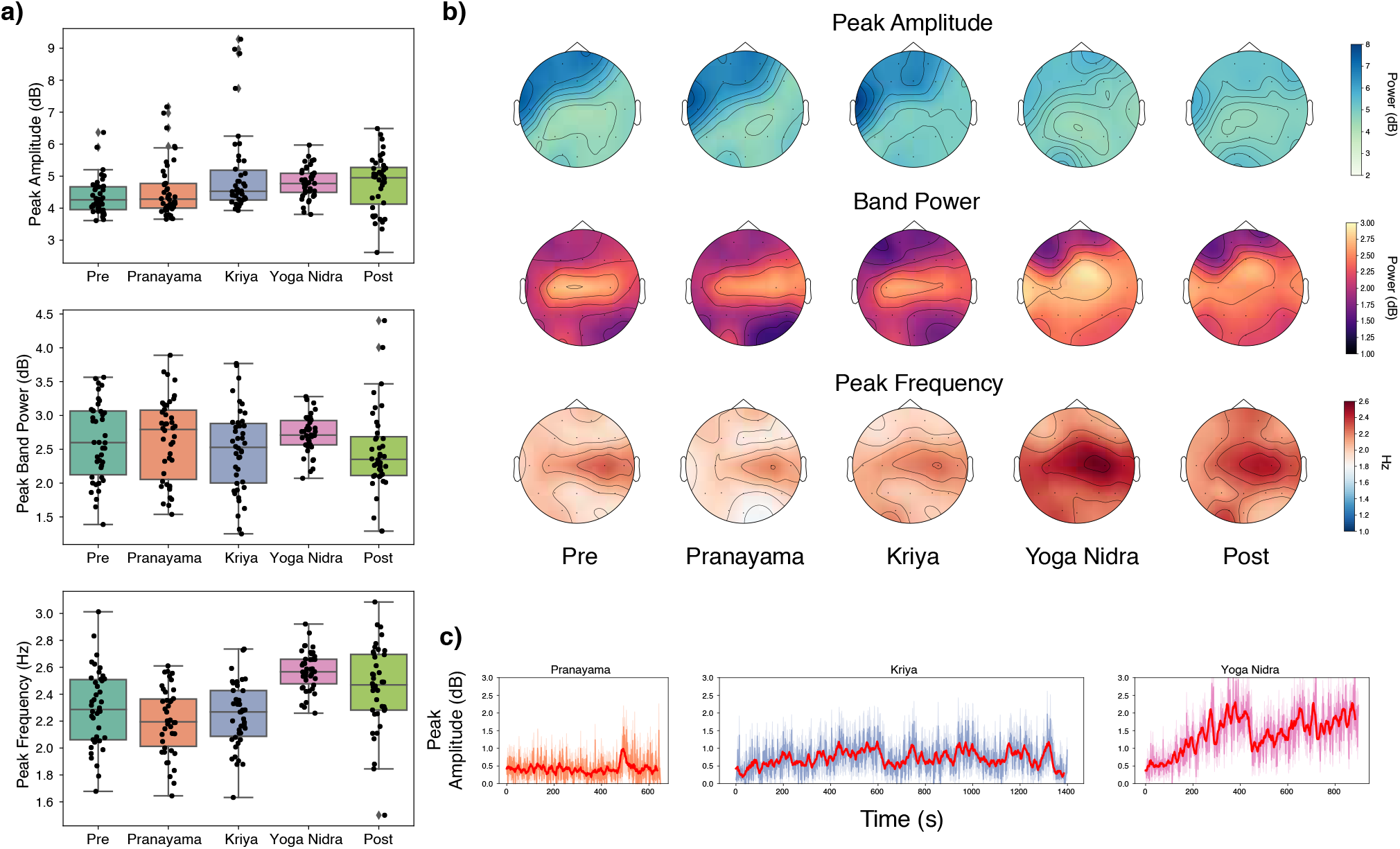
Effect on Delta wave dynamics: a) Bar plots show the peak amplitude, band power and frequency for the delta band computed using the FOOOF toolbox (Donogue et. al. 2020) for C4 electrode. b) Topomap plots for the peak amplitude, band power and peak frequency for the delta waves across the various parts of the SKY meditation. c) Change of peak amplitude across the meditation, we see a rapid increased in the delta amplitude during the the Yoga Nidra period.

**Figure 5.**
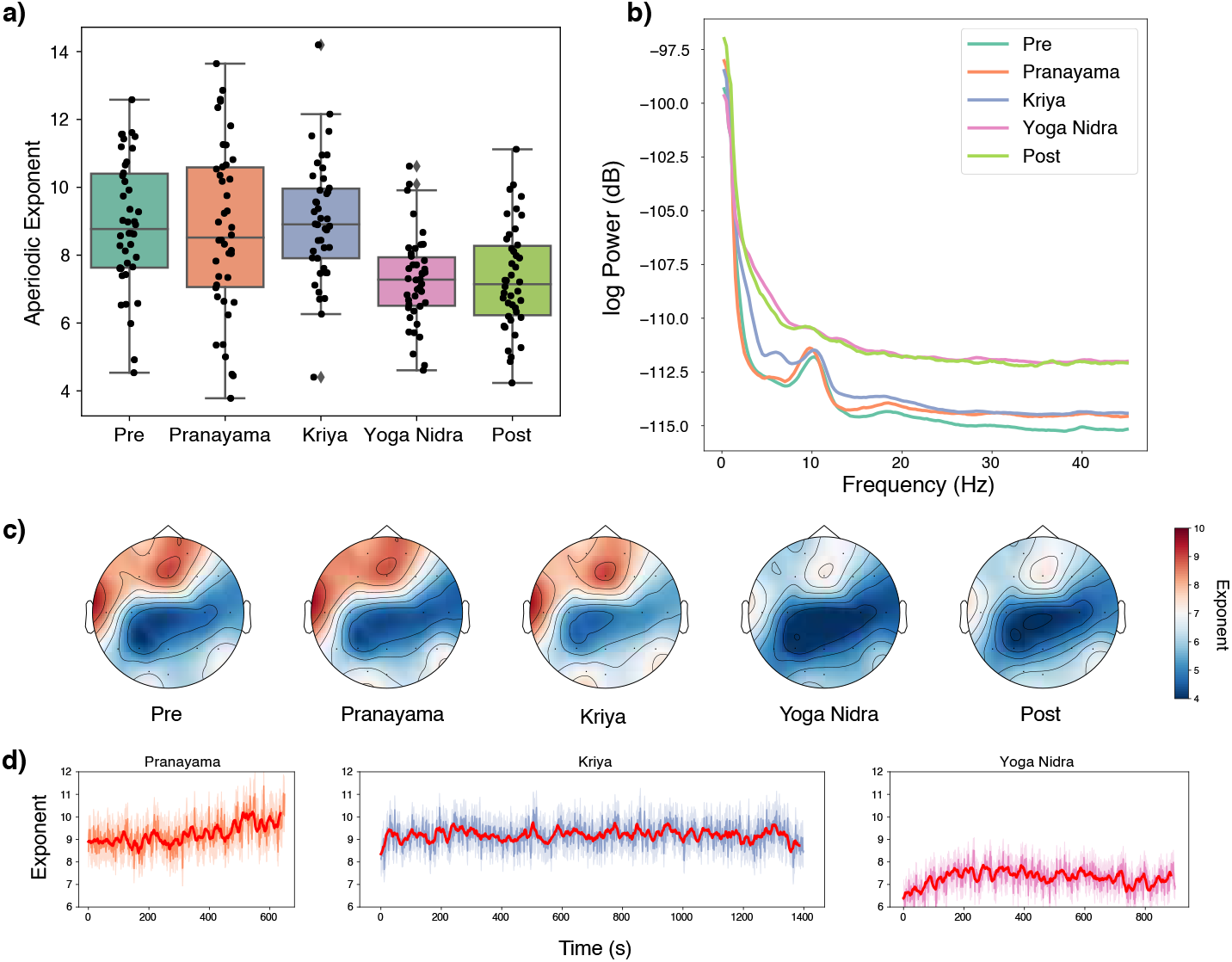
Effect on Delta wave dynamics: a) Bar plots show the peak amplitude, band power and frequency for the aperiodic 1/f component computed using the FOOOF toolbox (Donogue et. al. 2020) for C4 electrode. b) Frequency spectrum plots averaged across the five events. We see a decrease in the aperiodic 1/f slope during the Yoga Nidra and post resting phases. c) Topomap plots for the peak amplitude, band power and peak frequency for the aperiodic exponent across the various parts of the SKY meditation. d) Change of peak amplitude across the meditation, we see a rapid increased in the delta amplitude during the the Yoga Nidra period.

## Discussion

Breathing induces strong coherence in the neuronal oscillations as found using intracranial recordings in rodents and humans^10–14^. Respiration not only affects the neuronal activity in the olfactory systems and limbic systems but induces synchronized oscillations across the neocortex^13^ and higher brain circuits are more involved during conscious breathing than autonomic breathing^14^. The spatiotemporal dynamics of EEG during different phases of breath-based meditation were lacking and our research was aimed at filling that gap. We found that during the kriya period of the SKY meditation, the theta power increases and the yoga nidra period is associated with alpha power decrease, sustained theta, delta increase and decrease in the 1/f aperiodic signal exponent. The decrease in alpha and increase in delta/theta power during the meditation period is gradual but the reduction in the aperiodic exponent is sudden suggesting transition to deeper states of meditation.

Alpha power is associated with sensory feedback signal^36^ which propogates downstream the sensory hierarchy from the parietal to the sensory area and activates during periods of rest. The alpha amplitude in the parieto-temporal and occipital regions decrease as the SKY meditation progresses and we see a gradual decline in the amplitude during the yoga nidra period suggesting a decreased sensory feedback. But we see an increase in the alpha band power in these regions. A decrease in peak power and increase in the overall band power suggests a flattening of the alpha response in the brain. The decrease in peak alpha amplitude is consistent with earlier studies that the meditation phase is associated with decrease in the alpha amplitude^8,37^. And some other studies talked about increase in alpha power during meditation as compared to rest which could be related to the increase in alpha band power that we have observed^5,38–40^. 1/f exponent factor can bring in possible inconsistencies in alpha power computation as shown by Donogue and colleagues^33^. There could also be differences in meditation practices but researchers may also need to revise analysis based on the recent methods to remove the effect of aperiodic signals from band power computations. Alpha power is a characteristic frequency of the brain with relations with the Default Mode Network of the brain^41^. Reduction in the peak alpha amplitudes could also signify decrease in the Default Mode activity of the brain as studies have found reduced mind wandering during meditation^35^ and meditators have shown better anticorrelation with the Dorsal Attention Network which suggests better cognitive health^42^.

Human theta rhythm characterized by 4-7 Hz oscillations have been linked with various tasks like recognition^43^, spatial navigation^44–46^, working memory^43^, episodic memory retrieval^47^ and other complex cognitive tasks^48,49^. Theta rhythms possibly originate in the frontal midline regions including the prefrontal cortex and anterior cingulate cortex^50,51^ and have been associated with autonomic system^52^ and attentional engagement^53,54^. Various meditation practices like focussed attention, open monitoring, loving kinds and transcendental meditation have shown theta activity increase^7,55–57^ which is correlated with the training and expertise of the meditator. Increase in the frontal midline theta during meditation could suggest increase in the parasympathetic activity and internalized attention. We found that during the kriya breathing period the peak theta frequency increases in the fronto-central regions along with increase in the peak amplitudes but during the yoga nidra period, low theta peak frequency with an increased band power predominates which could be possibly due to the changes in the balance between sympathetic and parasympathetic nervous system related to rhythmic breathing.

We found that the peak delta amplitudes increase during the meditation period with an increase in the peak delta frequency in the fronto-central regions. The pranayama and kriya breathing periods did not invoke as sharp changes in the delta wave parameters. Some studies have shown a reduction in delta frequency^55^ during meditation but others^53^ have shown an increase in delta activity. Delta oscillations are involved in functions related to homeostasis and autonomic activity^58–60^. Sleep studies show that delta becomes prominent during NREM phases of sleep^61^. During the meditation period we see delta amplitudes in the range 2-3 Hz increase along with theta waves which could signify a state akin to sleep but with awareness which aligns with the definition of the state of *Turiya* according to Yogic texts.

Aperiodic or 1/f electrophysiological signals have been associated with different states of consciousness, cognitive tasks^62,63^ and aging^32^. Drug induced states of consciousness have specific 1/f signatures^34,63,64^. Recent development of methods^33^ have allowed us to estimate aperiodic signals from the frequency spectrum. Local field potential studies have found an inverse relationship between the excitation-inhibition (E:I) balance in the brain and the exponent of the aperiodic signal^30^. A recent meditation study showed that the aperiodic exponent increased during meditation^35^. In our study, we saw the decrease in the 1/f aperiodic signal during the meditation period suggesting an increase in the E:I balance. A study upon the administration of propofol found spectral exponents to increase (or decrease in E:I balance) which does align with our results suggesting different signatures for meditation and unconciousness. As meditations are different in their nature, it could be possible the aperiodic signature are different for various meditations as a recent study reported increase during meditation as compared to rest. Further research would be required to find out the reason for such a change.

We did not analyse the gamma and beta bands as change in the aperiodic signals could confound any changes in the gamma and beta band power and amplitude. Improved analysis techniques could help us eke our the role of gamma and beta frequencies in breathing based meditations.

The study analysed the dynamics during the breath-based meditation technique called Sudarshan Kriya Yoga (SKY) and we found that theta rhythms become prominent during the rhythmic breathing period with an increase in peak theta frequency and amplitude and the meditation period was marked with a decrease in peak alpha amplitude but increase in alpha band power, increase in low frequency theta amplitudes and band power. We found an increase in peak delta band power and frequency during the meditation period. We also saw a sharp decrease in the 1/f aperiodic exponent during the meditation phase suggesting a change in the excitation-inhibition balance of the brain. Ours is one of the first study to analyze the spatiotemporal dynamics of electrophysiological signatures during different phases of a breathing based meditation and we find that meditation influences the periodic as well as aperiodic activity in the brain and aperiodic exponent coupled with delta and theta frequencies could act as a bio-signature of meditation.

## Methods

### Experiment Design

#### Participants

We recruited forty three subjects (23 M, mean age = 25.45, S.D = 5.75) without a prolonged medication, any psychological disorder or with a history of epilepsy. Subjects were selected randomly from the Art of Living Foundation’s Centre in Bengaluru, Karnataka, India. The research was advertised by the Human Resources Department and interested volunteers were selected. The participants had learned the SKY meditation prior to our research and were regular practitioners. The experience with the SKY practice ranged from 1-18 years (mean=7 years). 15 subjects had more than 6 years of experience. Informed consent and permission was obtained from all subjects prior to the start of the study. Ethical Committee of Ved Vignana Maha Vidya Peeth provided the ethical clearance for the research.

##### Electroencephalography (EEG)

A 24-channel EEG system, Superspec 24 (Recorders and Medicare Systems, Chandigarh, India) was used for data acquisition. We placed the electrodes according to the international 10-20 system. EEG was acquired from the frontal (FP1, FP2, F3, F4, F7, and F8), centro-temporal (T3, T4, C3, C4, FZ, CZ, PZ, T5, and T6), parietal (P3 and P4), and occipital (O1 and O2) regions. Ground at the forehead and earlobe reference was used for all electrodes. The system had a sampling frequency of 256 Hz and the a band pass filter of 0.1-75 Hz was applied. Impedance levels were kept under 5 kΩ. A temperature of 25°C was maintained throughout the experiment and the light were kept dim to provide a comfortable space for meditation.

##### Procedure for Sudarshan Kriya Yoga

Sudarshan Kriya Yoga is a cyclic breathing and is done sitting with the spine erect. It is preceded by three-stage Pranayama (focussed breathing with posture) where the hands are kept in different positions for each stage. 6-8 *Ujjaayi* breaths (slight forceful breath with slight throat contraction) with a cadence of breath in -hold - breathe out - hold is maintained during the three stages. It is followed *Bhastrika* pranayama which involves forceful breaths in and out with movements of hand in up and down manner. Om chanting is done thrice followed the the Kriya rhythmic breathing which involves breathing in normal breath in three different rates (slow, medium and fast) as guided by a pre-recorded audio. It is followed by rest in supine position. Prior publications had desrcribed in detail the procedure for SKY^29,65^

##### Protocol

We recorded the EEG in supine position with eyes closed for five minutes before the SKY practice, during the SKY practice, and for five minutes after the SKY practice for a total of around 1 hr 15 mins. Audio guided instructions allowed consistency in the frequency of breathing for all participants.

### Analysis

#### Electroencephalogram analysis

We screened the data visually to remove mechanical and motion artifacts. F7 and O2 channels were removed due to excess noise across multiple subjects during the paradigm. We utilized the mono-polar montage of channels and removed the global average signal followed by re-referencing it with Cz. We used the Multitaper Spectral Analysis toolbox^66^ to compute the spectrogram. Frequency range was set between 0.1 and 45 Hz, time-half bandwidth of 4 seconds and number of tapers for the multitaper spectral estimation were set to 7. We used a moving window of 4 seconds with step size of 1 second which resulted in frequency spectrum at each second of the entire SKY technique. We then used the Fitting Oscillations and One Over F(FOOOF) toolbox^33^ to extract spectral parameters from spectrum at each time point resulting in number of values as the number of time points. The FOOOF toolbox extracts the aperiodic signal and then computes the peaks from the spectrum without any explicit definition of the band ranges resulting in a better estimate of the peak amplitude and band power. We characterized peaks corresponding to delta as between 1–4 Hz, theta between 4.5–7.5 Hz, and alpha between 8–13 Hz. Aperiodic signal had an exponent and offset component. We extracted the peak frequency, amplitude and band power for the three rhythms and the aperiodic exponent value at each spectrum. We then divided the entire duration of the run into five components: Resting pre, Pranayama, Kriya, Yoga Nidra and Resting post and averaged the peak frequency, amplitude and band power components across these events.

#### Statistical analysis

In order to compare the peak frequency, amplitude and band power across the different events, we performed a one-way repeated measures ANOVA followed by Tukey’s pairwise post hoc analysis using the statsmodels package in Python 3.7^67^. We repeated it across the seventeen channels and bonferroni corrected to reduce multiple comparisons error.

## Acknowledgements

The study was financially supported by Ved Vignan Maha Vidya Peeth (VVMVP, Bangalore).

## Author contributions statement

V.K designed the study and experiments, L.B and V.K conducted the experiments, V.T analysed the data and wrote the first draft. All authors reviewed the manuscript and contributed to the final draft.

